# Genetic evidence of a hybrid swarm between Alpine ibex and domestic goat

**DOI:** 10.1101/2023.12.22.572967

**Authors:** Alice Brambilla, Noel Zehnder, Bruno Bassano, Luca Rossi, Christine Grossen

**Author notes:** **Corresponding author:** Alice Brambilla.

## Abstract

Improving the understanding of the causes and effects of anthropogenic hybridization is fundamental to ensure the conservation of wild species, particularly in the case of hybridization between wild species and their domestic relatives. Knowledge is missing for many species also because of a lack of appropriate tools for hybrid identification. Here, coupling genotype and phenotype analysis, we carried out an extensive investigation of ongoing hybridization in Alpine ibex *Capra ibex*, a mountain ungulate of conservation concern from a genetic perspective. By genotyping at 63 diagnostic and 465 neutral SNPs 20 suspected hybrids and 126 Alpine ibex without suspicious phenotype, representing eight populations across a major part of the species distribution, we found evidence for ongoing hybridization between Alpine ibex and domestic goat. We identified different levels of hybridization including back crosses into both Alpine ibex and domestic goat. Our results suggest a lack of reproductive barriers between the two species and good survival and reproductive success of the hybrids. Hybridization was locally intense, alike a hybrid swarm, but not spread across the rest of the species distribution. Most of the hybrids were discovered in two locations in the North-West of Italy, while random sampling of individuals from different areas did not provide evidence of recent hybridization. Our method, based on Amplicon sequencing of 63 diagnostic SNPs specifically developed for this purpose, allowed us to identify hybrids and back crosses up to the 4^th^-5^th^ generation and was suitable for genetic samples of different quality, although with varying levels of certainty regarding the exact number of generations passed since hybridization. Based on the paired analysis of genotype and phenotype we provide guidelines for a first identification of hybrids in the field and suggest a procedure for the reliable identification of hybrids.

## Introduction

In the last decades, the number of anthropogenic hybridization events, defined as hybridisation as a result of human action (McFarlane & Pemberton, 2019), increased because of human activity such as translocations of organisms outside their natural range and habitat modifications (e.g., Allendorf et al., 2001; Iacolina et al., 2019; Moroni et al., 2022; Negri et al., 2013). It is generally acknowledged that hybridization can play an important role in evolution (Genovart, 2009). It can enhance adaptations, as for example in the case of mountain hare *Lepus timidus* and brown hare *Lepus europaeus*, where higher levels of introgression from the local species (mountain hare) into the expanding species (brown hare) were observed (Levänen et al., 2018). By acquiring functional variation from the local mountain hare, brown hare may be adapting more easily to the new environment (Pohjoismäki et al., 2021). Hybridisation can even prevent the extinction of endangered populations by genetic rescue (Pimm et al. 2006). Nevertheless, anthropogenic hybridization is widely recognized as a threat for conservation (Rhymer & Simberloff, 1996; Simberloff, 1996) as it may jeopardize the long-term genetic conservation, lead to the loss of local adaptations (Allendorf et al., 2001), cause genetic swamping (Howard-McCombe et al., 2023; Senn et al., 2019) or even cause the local extinction of wild species (Rhymer & Simberloff, 1996). Even more unanimously, anthropogenic hybridization between wild species and their domestic relatives is considered detrimental for the conservation of the wild taxon (Allendorf et al., 2001; Randi, 2008; Senn et al., 2019). Interestingly, although introgression of domestic genes may result in outbreeding depression and reduced fitness of hybrids (Fukui et al., 2018), it may also lead to higher reproductive success (e.g., hybrid vigor, Ferguson et al., 1988) and to the enrichment of some genes, as for instance, immune-relevant genes in Scottish wildcat *Felix silvestris* (Howard-McCombe et al., 2023), in Alpine ibex *Capra ibex* (Münger et al., 2023) as was previously shown also for archaic introgression into modern humans (Abi-Rached et al., 2011;Gouy et al., 2020). Hence, introgression may be adaptive under certain circumstances, particularly in species with low genetic diversity, increasing the chances of introgression to occur and thereby, at the same time, increasing the risk of genetic swamping (Howard-McCombe et al., 2023). The probability, spreading and consequences of successful hybridization, as well as the chances of back-crossing and hence of introgression to occur depend on several factors, among which the ecology and life history of the species of interest. However, most of the knowledge on the consequences of anthropogenic hybridization on the genetics of wild vertebrates comes from studies conducted on fish (e.g., Harbicht et al., 2014; Skaala et al., 2019). In the case of mammals, not many cases of hybridization, especially ongoing hybridisation, between wild species and their domestic relatives have been studied in detail from a genetic perspective and different effects were reported for different taxa (Adavoudi & Pilot, 2022). For instance, local introgression from domestic pig *Sus scrofa domestica* have been observed in wild boar *S. scrofa* populations in Europe (Canu et al., 2016). Management and high harvesting rates of wild boar have been suggested as potentially favouring the spread of the domestic genes but causes and consequences of introgression are not yet clear (Dzialuk et al., 2018). In the case of grey wolf *Canis lupus*, Pilot et al. (2018) suggested that hybridization with domestic dog *C. lupus familiaris* occurred repeatedly over time in different parts of Europe and Asia but, despite that, genetic differentiation between wolf and domestic dog has been maintained. On the contrary, hybridization with domestic cat is considered as the major threat for the critically endangered population of wild cat in Scotland (Senn et al., 2019). In addition, although hybridization with the domestic cat enriched immune-related genes in wild cats, allowing them to improve the immune response against diseases spread by domestic cat, maladaptive genetic swamping was associated with such enrichment (Howard-McCombe et al., 2023). The species-specific likelihood of occurrence and consequences of hybridization, paired with a lack of knowledge for most of the species, constitutes an obstacle to the implementation of effective and informed management actions for conservation. Filling this gap of knowledge is therefore pivotal to favour the conservation of genomic diversity of wild species, which is one of the objectives included in the Kunming-Montreal Global Biodiversity Framework (GBF) approved in 2022 during the fifteenth meeting of the Conference of the Parties (COP 15). Guidelines improving the identification of hybrids could constitute a first tool to foster research on hybridisation.

The Alpine ibex (*C. ibex*) is a mountain ungulate of conservation concern. Despite it being listed as LC in the IUCN Red List of species (Toïgo et al., 2020), its recent history makes it worth special attention from a genetic perspective: genetic variation of Alpine ibex is among the lowest observed in wild mammal species (Biebach & Keller, 2010; Grossen et al., 2020) as a consequence of the strong and repeated bottlenecks that occurred in the 19th century (Stüwe & Nievergelt, 1991) increasing inbreeding and inbreeding depression (Bozzuto et al., 2019; Brambilla et al., 2014). Because of its low genetic diversity, high levels of inbreeding and relatively high genetic load (Biebach & Keller, 2012; Grossen et al., 2018, 2020), Alpine ibex may be particularly susceptible to successful hybridization leading to introgression. As the genus *Capra* is relatively young, all *Capra* species can interbreed and produce fertile offspring (Couturier, 1962; Pidancier et al., 2006). The domestic goat (*Capra hircus*) is the only *Capra* species occurring in the same region as free-ranging Alpine ibex and hybridization has been reported both in captivity and in the wild (Iacolina et al., 2019).

A successful hybridization event between Alpine ibex and domestic goats happened in the recent past and lead to introgression still visible at the MHC region (Grossen et al., 2014). This event probably occurred during the bottleneck of 18^th^-19^th^ century when the Alpine ibex population was at its minimum, confined in the Gran Paradiso region (North-Western Italian Alps). Indeed, the Gran Paradiso area was home of the only remnant population from which all the reintroduced Alpine ibex populations in Europe ultimately derived. Afterwards, the likely adaptive MHC alleles increased in frequency and spread with the reintroductions across the Alps (Grossen et al., 2014).

Less was known, instead, on ongoing hybridization in Alpine ibex. It has sporadically been reported (Giacometti et al., 2004; Iacolina et al., 2019) but no quantitative data about the extent of the phenomenon were available until recently, when a large-scale survey conducted across the Alps allowed to gather systematic information on the presence of suspected hybrids (Moroni et al., 2022). The study revealed that hybrids were actually present in most of the European countries with at least 48 probable hybrids observed between 2000 and 2021. However, the study identified probable hybrids based on phenotypic characteristics while no genetic confirmation was available for most of the animals. The same work (Moroni et al., 2022) also highlighted the lack of knowledge about the potential effects of ongoing hybridization on the genetic structure of the species (e.g., whether hybrids were able to further reproduce with Alpine ibex enabling introgression of domestic goat alleles into Alpine ibex populations).

### Aims

The aim of this work is to shed light on the extent and the genetic characteristics of hybridization between Alpine ibex and domestic goat and to understand whether phenotypic appearance could be a reliable tool to assess hybridization. To do so, we analysed samples of suspected and non-suspected hybrids using a genomic diagnostic tool specifically developed for this species pair (Kessler et al., 2022). Samples were collected in two areas identified as hotspots of potential hybridization by Moroni et al. (2022) as well as in other areas where no suspected hybrids were observed. In detail: 1) we assessed the hybridization status and ancestry of individuals that were identified as suspected hybrids based on their phenotypic characteristics; 2) we screened the rest of the target populations as well as other populations where no suspected hybrids were reported to find possible hybrids that were not identified from phenotypic characteristics; and, finally, based on the obtained results, 3) we provided recommendations for the identification of hybrids and suggested further application of the method to other species.

## Methods

### Sampling

A total of 159 samples were used for this study: N=20 from suspected hybrids, N=126 from individuals with no phenotypic signs of hybridization (non-suspected individuals) and N=13 from domestic goats (Figure 1A). More details on provenience, collection, and handling of the different kind of samples are provided in the next paragraphs.

**Figure 1:**
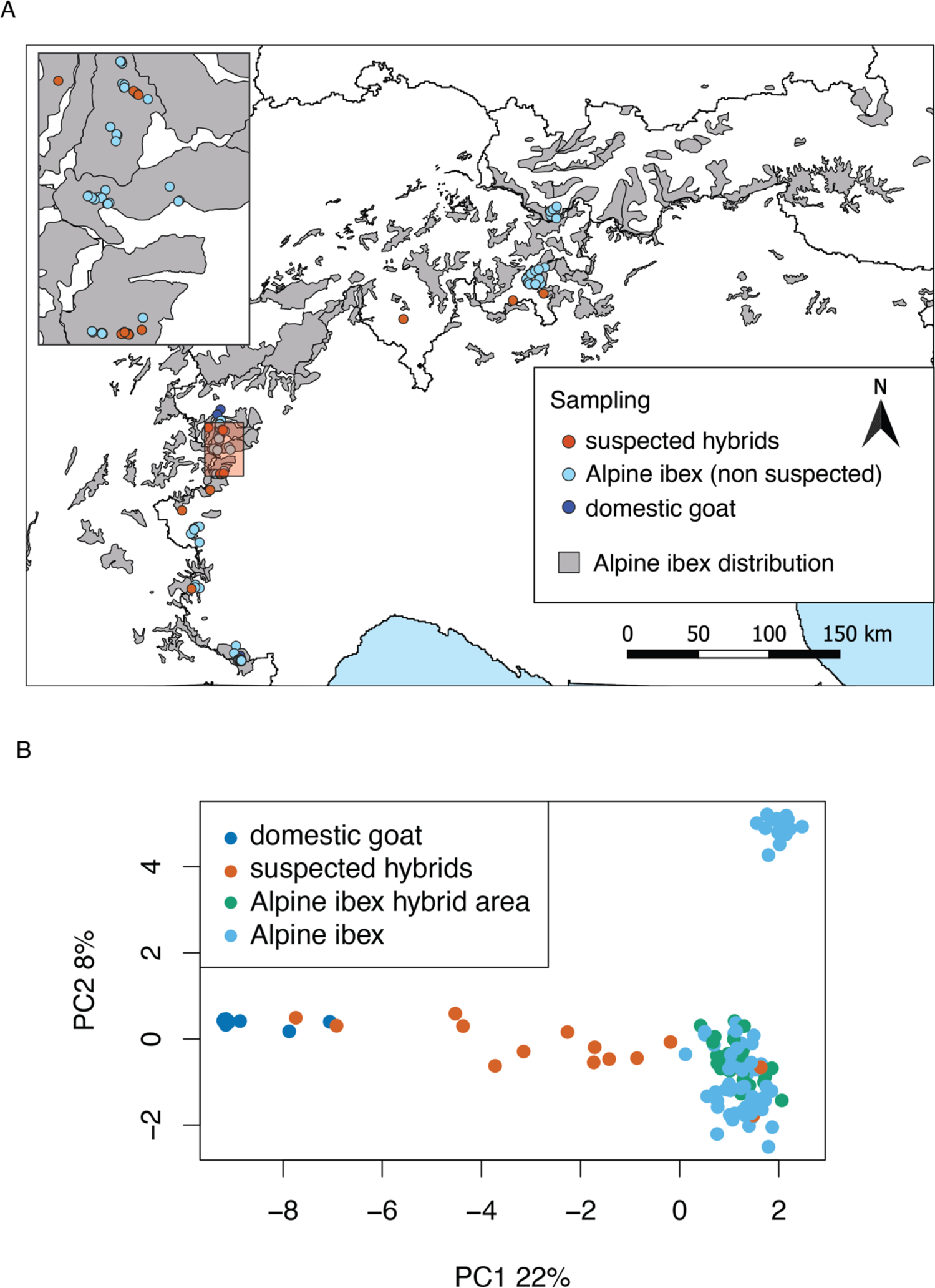
A) Map of the location of the samples included in the study. In orange the suspected hybrids, in light blue the non-suspected Alpine ibex (collected both in the focal areas – where several suspected hybrids were observed - as well as in other areas where no suspected hybrids were observed) and in dark blue the domestic goats used as reference (non-suspected to be hybrids). B) Principal component analysis (PCA) based on 465 neutral SNPs on 133 individuals: (light green: non-suspected Alpine ibex outside focal area (Alpine ibex hybrid area), light blue: non-suspected Alpine ibex focal area (Alpine ibex), orange: suspected hybrids, and dark blue: non-suspected domestic goats (domestic goat). See Figure S1 for the same graph including population information.

### Suspected hybrids

In total we analysed 20 samples from suspected hybrids (see Supplementary Table S2). 13 samples (from 12 suspected individuals) were specifically collected for this study between 2020 and 2022 in two populations identified as hotspots of potential hybridization by Moroni et al. (2022): Lanzo (Val d’Ala di Stura, TO, Italy, N=10) and Gran Paradiso (Valsavarenche-Rhemes, AO, Italy, N=3). In addition, samples from 7 suspected hybrids collected in the past years in different parts of the Alps were opportunistically added to the sample set: Ticino (TI, N=1), Graubunden (GR, N=2) and Fribourg (FR, N=1) in Switzerland and Val di Susa (TO, N=2) and Val Varaita (CN, N=1) in Italy (see also Figure 1A).

The specimens included tissue, blood and faecal samples. Tissue and blood samples were collected during captures (N=4 samples in Lanzo), after culling of the suspected hybrids (N=3 samples in Gran Paradiso and N=3 in Switzerland: N=2 in Graubunden and N=1 in Fribourg), or in stable from domestic individuals suspected to be hybrids (N=1 in Lanzo, N=1 in Ticino and N=1 in Val di Susa). Faecal samples (N=7 from Lanzo) were collected in the field: suspected individuals were identified and followed from distance. Immediately after defecation fresh pellets were collected in plastic bags and frozen in liquid nitrogen. One of the faecal samples came from one of the captured individuals, to allow comparison of genotypes obtained from different kind of samples.

### Non-suspected individuals

To test whether phenotypic appearance was a reliable tool for hybrid identification and to conduct a genetic screening of populations where no suspected hybrids were reported, we also analysed 126 samples (N=88 tissue or blood and N=38 faecal samples) from individuals with no phenotypic signs of hybridization collected in the two areas with hybrid presence (Valsavarenche, N=26 (Gran Paradiso population, sympatric with the observed suspected hybrids); Lanzo, N=16 (Lanzo population, sympatric with the observed suspected hybrids) as well as in six additional, supposedly hybrid-free populations or sub-populations: Valle Orco, N=14 (TO, Italy, Gran Paradiso population but geographically distant from the observed hybrids); Lanzo, N=12 (TO, Italy, Lanzo population but distant from the observed hybrids); Albris, N=19 (GR, Switzerland); Alpi Marittime, N=16 (CN, Italy); Queyras, N=15 (between Italy and France); Silvretta, N=12 (Austria).

Captures and culling were conducted trying to minimize adverse effects on animal welfare and were authorized by the local authorities. Captures in Gran Paradiso were authorised by the Italian Ministry of the Environment (authorisation no. 25114 of 21/09/2004), periodically reviewed by the Italian National Institute for Environmental Protection and Research. Captures in Lanzo were authorized by ASL TO4 and Città Metropolitana di Torino.

### Domestic goats

Finally, to have a comparison with domestic goat breeds typical of the sampling areas (i.e., the potential sources of hybridization), we also analysed 13 tissue samples collected from domestic goats belonging to local herds (N=2 from Lanzo, N=6 from Gran Paradiso, N=5 from Alpi Marittime region).

### Phenotypic data

All the suspected hybrids sampled for this study were photographed and information about sex, age and deviations from the common phenotypic appearance of Alpine ibex were collected. After identifying the most common phenotypical anomalies of the suspected hybrids (i.e., diagnostic traits) we summarized them in the following four categories, which were then evaluated for each individual: horn (anomalies in horn shape, size, presence/absence), pelage (anomalies in pelage colour or length), muzzle (muzzle and front appearance more similar to those of domestic goats), body (body size or shape different from that of Alpine ibex, but also presence of visible udders in females or very large testicles in males). Non suspected Alpine ibex included in the dataset were also screened for the same diagnostic traits.

### DNA extraction

Blood samples were collected in EDTA vacutainer and stored at +4°C (with the exception of a sample that was conserved on an FTA© card), tissue samples were stored at room temperature in ethanol 97-99%, and faecal samples were frozen in liquid nitrogen immediately after collection and subsequently stored at −20°C until analysis. DNA extractions from the faecal samples were performed using the QIAamp DNA Stool Mini Kit (QIAGEN) while for blood and tissue samples the DNEasy Blood & Tissue Kit was used.

### Genetic data generation

All samples were genotyped using a microfluidics-based amplicon sequencing assay (Juno system, Fluidigm) designed by Kessler et al. (2022). The assay was designed to cover 1’265 amplicons and includes 84 diagnostic markers for Alpine ibex and domestic goat, 744 neutral markers and 433 MHC and other immune-related markers (the latter not used for this study).

Sample preparation and amplicon amplification was performed according to the manufacturer’s protocol except for the amount of input DNA (concentration normalized to a maximum of 50 ng/µl, lower where this concentration could not be reached), see Kessler et al. (2022) for more details. Sequencing of the libraries was performed on a NextSeq 500 system (Illumina, San Diego Ca, USA) in mid-output mode. To avoid potential problems due to low sequencing complexity, 30% PhiX was added. Amplification and sequencing were repeated two to three times for all faecal samples while a single analysis was done for tissue and blood samples. Sample processing up to sequencing were performed in the framework of a larger monitoring project and hence jointly with samples which are not part of this study.

The sequenced reads were demultiplexed with default settings using Bcl2fastq v2.19 (Illumina®, 2019) and trimmed with Trimmomatic v0.36 (Bolger et al., 2014). Forward and reverse reads were merged using Flash v1.2.11 (Magoč & Salzberg, 2011) and then mapped to the domestic goat reference genome ARS1 (Bickhart et al., 2017) using bwa mem 0.7.13 (Li, 2013). The mpileup and call functions of Bcftools v1.17 (https://www.htslib.org/) were used for variant calling. Bcftools mpileup was run with min-MQ 10, only retaining sites with a minimal mapping quality of 10 and -a DP, AD, ADF, ADR to retain read depth information. Bcftools call was run using -mv -Ob -f GQ for outputting only variant sites in default calling mode, outputting to bcf-format and outputting genotype qualities. Repeats of faecal samples were trimmed, merged and mapped separately, but to combine all repeats per individual into one genotype per individual at the genotyping stage, the individual ID was used as read group (RG) identifier in bwa.

Vcftools (Danecek et al., 2011) was used to convert to vcf format, set the minimal genotype quality to 50 and the minimal read depth to 20. In the next step, we removed individuals with high missingness rate (>80%) and only kept bi-allelic SNPs with a genotype rate of at least 80% over the retained individuals.

### Data analysis

For a first uninformed visualization of the samples, a PCA was performed based on the putatively neutral markers (133 individuals and 465 sites) using the R packages (R Core Team, 2022) {adegenet} and {vcfR}.

Next, to specifically test which suspected hybrids were genetically confirmed to be hybrids, for each individual we calculated the proportion of domestic goat ancestry based on the diagnostic markers (63 sites retained after filtering, distributed across all autosomes). The proportion of domestic goat ancestry of an individual was calculated by dividing the number of alleles known to be private for domestic goats based on the diagnostic markers by the total number of alleles genotyped at the diagnostic markers. An individual was defined as confirmed hybrid if at the diagnostic markers more than one domestic goat allele was observed. This was to avoid the erroneous classification as hybrid due to potential genotyping errors and also because only one out of 126 possible goat alleles (in 63 diagnostic markers) would mean hybridisation going back a very large number of generations (at least 6), which we would not anymore consider as a recent hybrid.

While the proportion of domestic goat alleles can give some indication on how many generations back the hybridisation event happened, genotype frequencies can be used to predict this with more precision. We used the tool NewHybrids (Anderson, 2003), which is based on the statistical framework by Anderson & Thompson (2002) to infer the posterior probability of an individual being of a specific hybrid category. We used a custom-made genotype frequencies file as input to allow the categorisation of up to fourth generation hybrids with different back-crossing possibilities (See Supplementary Table S1). PGDSpider v 2.1.1.5 (Lischer & Excoffier, 2012) was used to convert vcf to the input format of NewHybrids. After rerunning NewHybrids starting from different seeds and checking for the robustness of the outcome, we ran it with seeds unset (--s 0 0, seed defined by clock) for a burnin of 20’000 (--burn-in 20’000) followed by 100’000 replicates (--num-sweeps 100’000), with PiPrior=JEFFREYS and ThetaPrior=JEFFREYS.

NewHybrids provides an objective approach to hybrid class identification. However, due to similar expected genotype frequencies, certain hybrid classes can still not be distinguished purely based on observed genotype frequencies, because potential recombination events are not accounted for. Thus, we carried out visual inspection of the genotypes. We created “chromosomal paintings” along the 29 autosomes to visualize the genotypes at the 63 diagnostic markers for all confirmed hybrids (i.e., individuals presenting more than one domestic goat alleles).

For an explorative comparison between different kind of samples, we repeated the same analysis (calculation of proportion of goat alleles, NewHybrid and chromosomal painting) on a tissue and a faecal sample collected from the same individual.

We finally tested the reliability of the phenotypic traits as diagnostic tool for the identification of the hybrids. Four phenotypic traits (hereafter: diagnostic traits): horn, pelage, muzzle, body, were identified as being potentially diagnostic as they deviated from the common Alpine ibex phenotype in the suspected hybrids. For each individual, we counted the number of diagnostic traits which ranged between 0 (no traits) and 4 (anomalies in all the identified traits). In addition, each animal was assigned to one of the following categories: no diagnostic traits (none of the diagnostic traits was noticeable in the animal), horn only, pelage only, muzzle only, body only and more diagnostic traits (more than one diagnostic trait was present). Pure domestic goats were excluded from these analyses.

A logistic regression was then built, where the probability of being a hybrid (binary variable with 0 for non-hybrid Alpine ibex and 1 for genetically confirmed hybrid, i.e., those individuals with more than one domestic goat allele, see previous paragraph for the rationale behind it) was dependent from the number of diagnostic traits. Furthermore, a binomial model was built with the proportion of domestic goat alleles being dependent from the number of diagnostic traits.

Because of the low number of observations of animals with only one diagnostic trait (pelage only N=3, horn only N=1, body and muzzle only=0) it was not possible to test the reliability of specific single traits as being diagnostic for hybridization. Graphical representation of the raw data was then produced.

## Results

After filtering for minimal genotype quality, read depth and genotyping rate, 133 samples out of 159 were retained and genotyped at 1226 sites. Of the 26 samples not passing the missingness filter, the majority (N=19) were faecal samples, six were tissue samples mostly collected from animals found dead (hence likely degraded DNA) (Supplementary Figure S1) and one was blood drops on filter paper (FTA© card). The sample with highest missingness retained in our analyses had 16 diagnostic marker genotypes, none of which suggested hybridisation. Among the 133 samples passing the filters, 17 belonged to suspected hybrids (representing 16 individuals because of the replicate sampling of one of them, see below). Of the 1226 sites, which also include MHC-linked and other immune-related markers (Kessler et al., 2022), 465 neutral and 63 sites were used for this study (see more details below). Median individual read depth was 151 and ranged from 11 to 354.

A first visualisation of the 133 samples analysed in this study using a principal component analysis (465 neutral sites) clearly separated Alpine ibex from domestic goat (Figure 1B, PC1, 22% variance explained). All Alpine ibex clustered together except for the population of Alpi Marittime (Supplementary Figure S2, PC2, 8% variance explained), a population which was previously shown to be genetically distinct from all other Alpine ibex populations (Grossen et al., 2020; Kessler et al., 2022). The majority of the suspected hybrids were placed along a continuum between Alpine ibex and domestic goat, while two suspected hybrid individuals clustered with domestic goat and two others with Alpine ibex (Figure 1B). Domestic goat clustered more closely together than Alpine ibex not because they are less diverse than ibex, but because the amplicon assays used here were designed using Alpine ibex as a reference and hence an ascertainment bias is present (see also Kessler et al., 2022).

The specific analysis of the 63 diagnostic markers revealed a wide spectrum of proportion of domestic goat ancestry for the suspected hybrids. All suspected individuals based on phenotype, except for four individuals which only showed a slightly unusual fur colouring as diagnostic trait (see below), were confirmed to be hybrids by the genetic analysis (Figure 2A, Supplementary Figure S3) for a total of 12 confirmed hybrids among the 16 suspected (6 males and 6 females, ages ranging from 2 to more than 15 years old). All 121 non suspected ibex and all the 13 domestic goats without any anomalies among the four phenotypic traits analysed were confirmed to be non-hybrids (proportion domestic goat equal to 0 or 1 respectively). The two non-suspected hybrids that had proportion domestic goat >0 but did not show any phenotypic diagnostic traits had a very low proportion of domestic goat alleles (≈ 0.01) and were genotyped from faecal samples, so we cannot exclude genotyping errors (see discussion). In addition, they were very young (a female of 2 and a male of 3 years old), hence, even if present, phenotypic diagnostic traits may still have been hidden.

**Figure 2:**
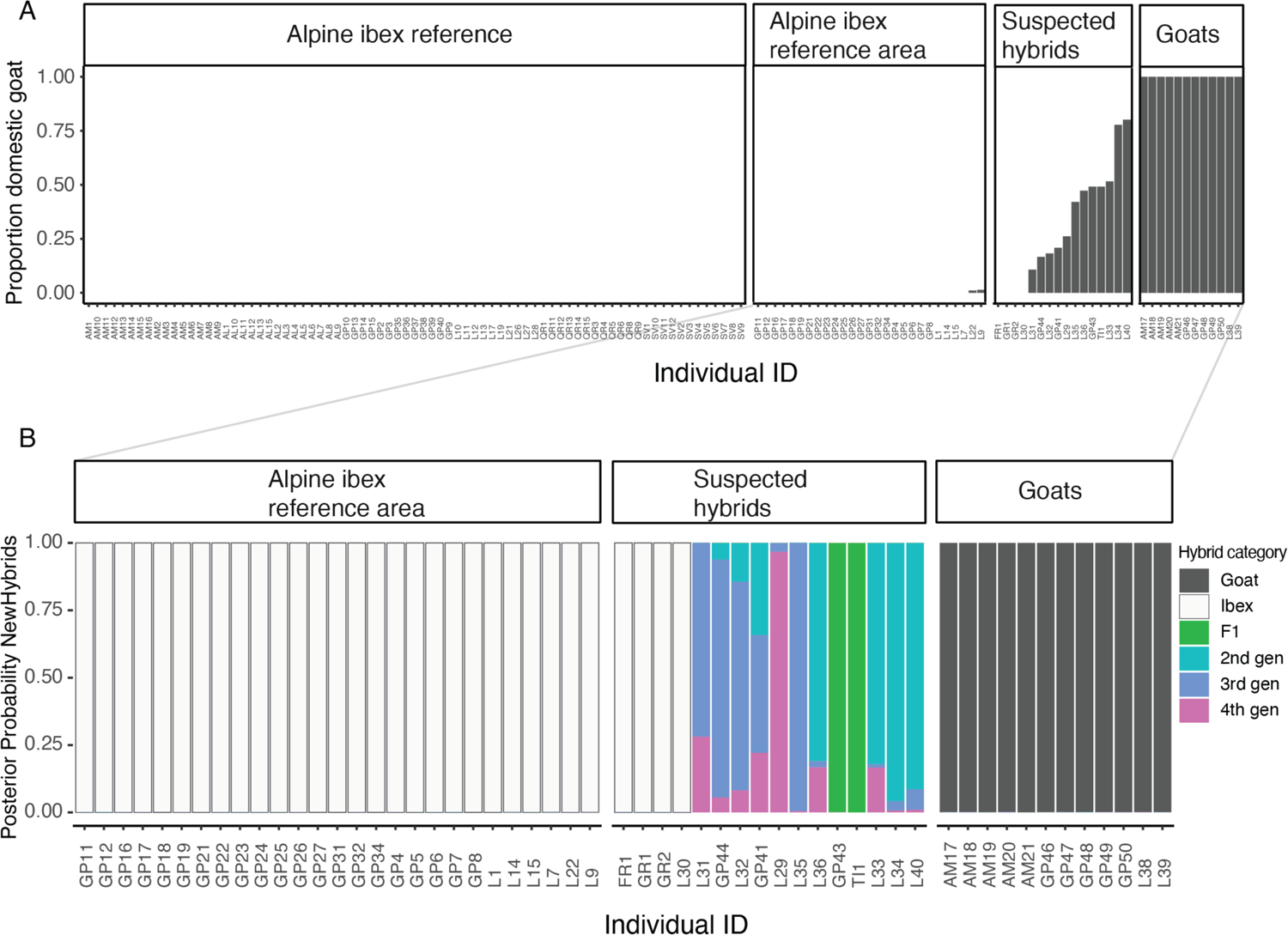
A) Proportion domestic goat ancestry based on 63 diagnostic SNPs, shown in bars (see Supplementary Figure S3 for the same figure including individual missingness). B) Cumulative probabilities computed by NewHybrids for each individual to belong to one of the pre-defined hybrid classes. The first-generation category F1 is a cross between an Alpine ibex (Ibex) and a domestic goat (Goat). Second-generation crosses (2nd gen) include F2 (Cross between two F1 hybrids), BxG (backcross of F1 with goat) and BxI (backcross of F1 with Alpine ibex). The third- and fourth-generation backcrosses (3rd and 4th gen) comprise all cross possibilities between second, first and “pure” generation (Alpine ibex or domestic goat) individuals. (See also Supplementary Figure S4 for a visualization of all tested categories). Individual IDs contain abbreviated population information: GP (Gran Paradiso), L (Lanzo), AL (Albris), AM (Alpi Marittime), QR (Queyras), SV (Silvretta), FR (Fribourg), TI (Ticino), GR (Graubunden).

The analysis with NewHybrids allowed the objective classification of the hybrids into hybrid classes (for instance, F1, F2, backcross with Alpine ibex). It confirmed what the analysis based on the proportion of domestic goat alleles suggested: among the hybrids, there were back-crosses both into Alpine ibex as well as into domestic goat up to 3^rd^ generation hybrids (Figure 2B), as expected in a “hybrid swarm”. Back-crossing into Alpine ibex and domestic goat was also shown by the additional investigations along chromosomes, which also revealed possible recombination events (Figures 3 and S4). The explorative comparison of the two sample kinds (tissue and faecal) from the same animal showed that hybridisation was clearly confirmed, and the individual was classified as third-generation hybrid in both cases. (Figure 4). However, the proportion of goat alleles as well as the exact hybrid class diverged between the two samples (Figures 4, B, C, and Supplementary Figure S5).

**Figure 3:**
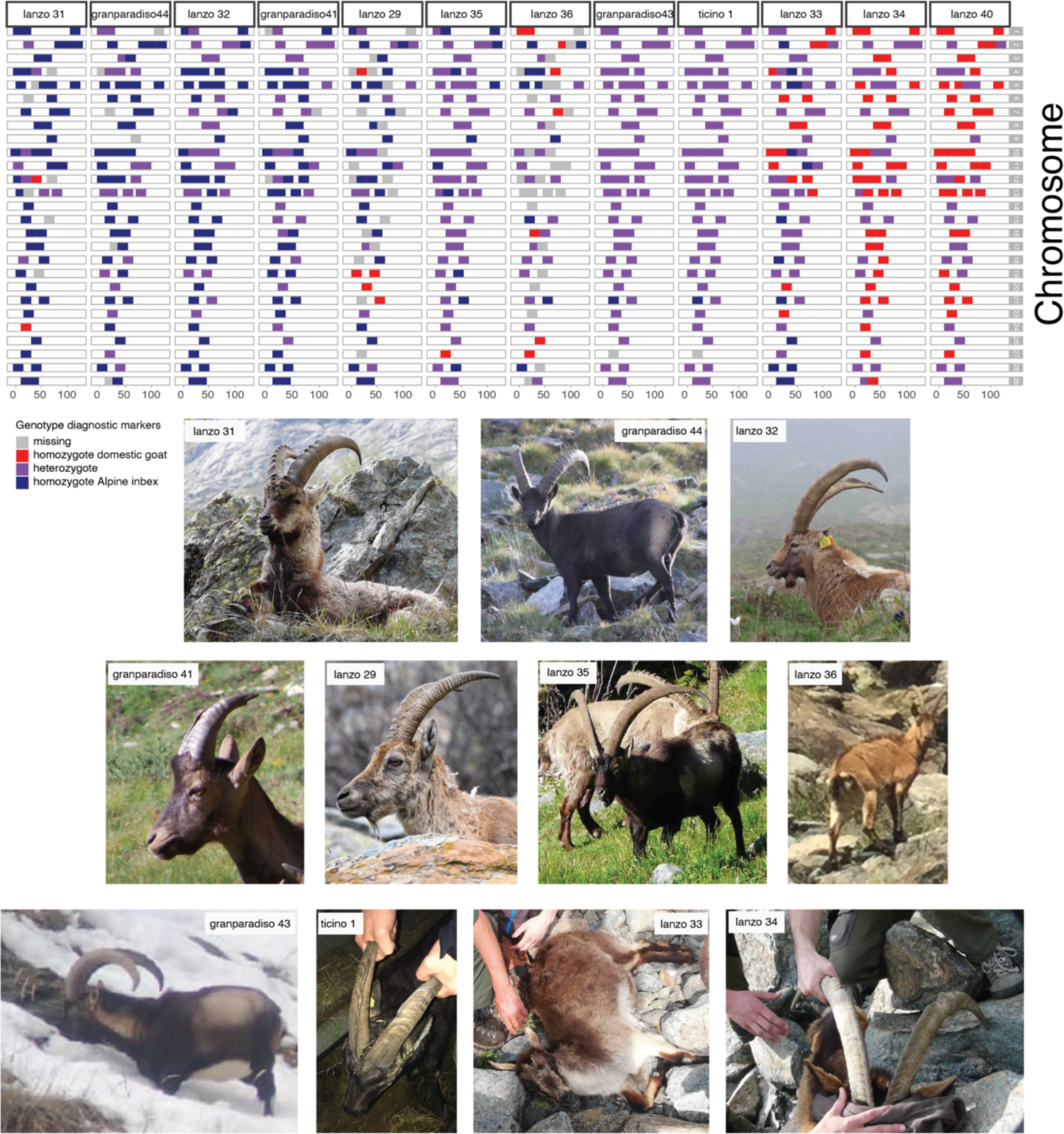
Chromosome painting with photographs: Chromosome painting along the 29 autosomes showing the diagnostic marker genotypes for all suspected hybrid individuals with a proportion of domestic goat alleles of at least 5% ordered by domestic goat proportion. Colours indicate heterozygous (purple), domestic goat (red), Alpine ibex (dark blue) as well as missing (grey) genotypes. Lanzo 40 was a domestic animal suspected to be hybrid (picture missing).

**Figure 4:**
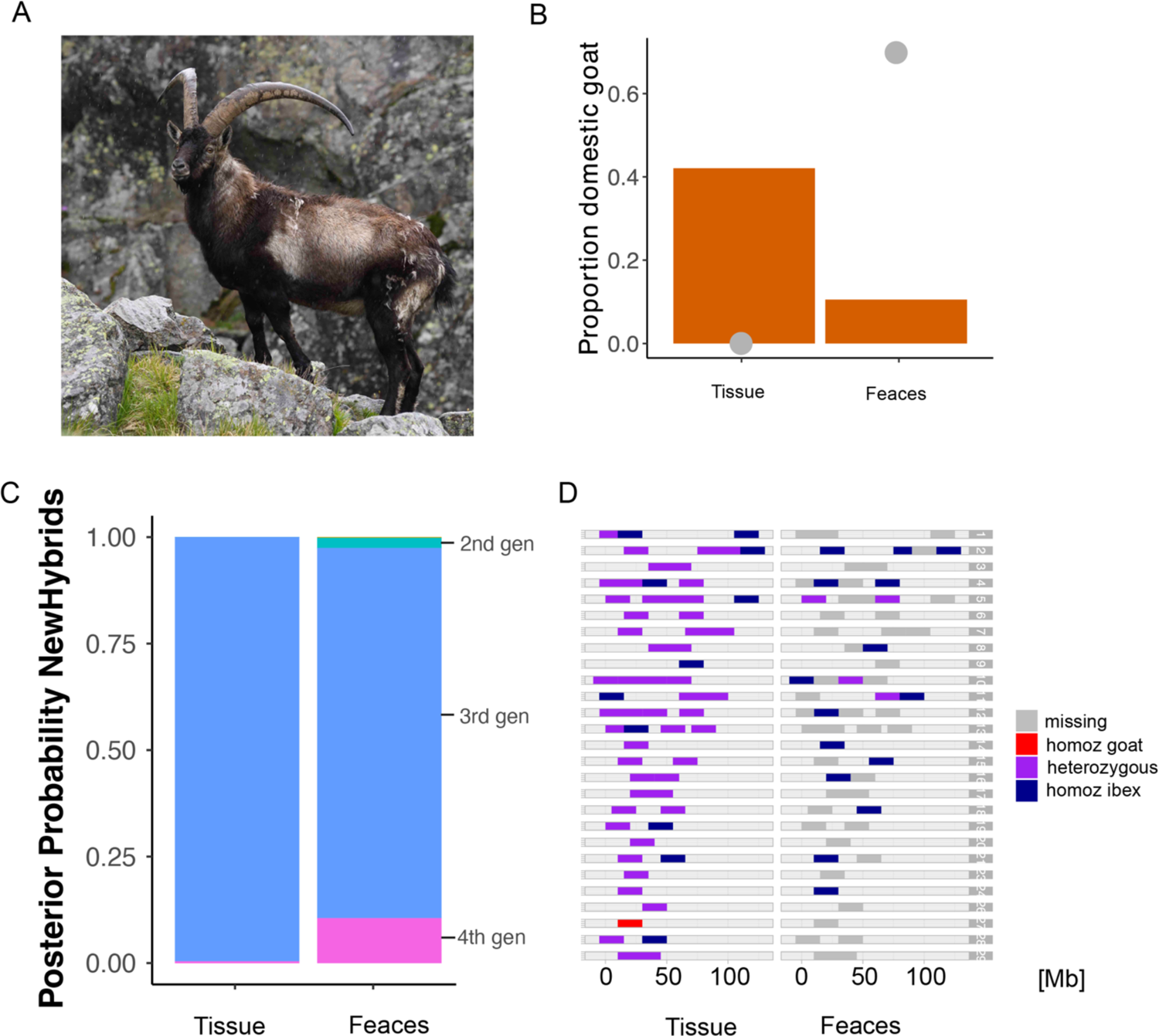
Comparison of results of different analysis to assess hybridization carried out on a tissue and a faecal sample from the same individual. A) picture of the individual. B) Proportion of domestic goat ancestry in the tissue and in the faecal sample. Grey dots represent missing loci (N=0 for tissue sample and N=44 (69.8%) for faecal sample). C) Cumulative probabilities computed by NewHybrids to belong to one of the pre-defined hybrid classes for the two samples (see also Supplementary Figure S5 for a visualization of all tested categories). D) Chromosome painting along the 29 autosomes showing the diagnostic marker genotypes for the two kinds of sample.

The proportions of hybrid and non-hybrid individuals that presented different diagnostic phenotypic traits are provided in Figure 5A. Most (11) of the 12 confirmed hybrids presented more than one diagnostic trait but one of them had only anomalies in the horns. Nearly all the non-hybrid Alpine ibex (N=104, 96.3%) had instead no phenotypic anomalies with the exception of three individuals in Switzerland that had the pelage of a colour lighter than the common Alpine ibex and a female older than 15 years old in Lanzo which was classified for two diagnostic traits (lighter pelage as well and particularly long horns, which anyway could be due to the very old age of the female) but were not confirmed as being hybrids by genetic analysis. It is worth to be noted that those individuals were considered as suspected hybrids in this study (because they showed at least one diagnostic trait) but none of them was confirmed as probable hybrid by the authors of Moroni et al. (2022) nor from our genetic analysis.

**Figure 5:**
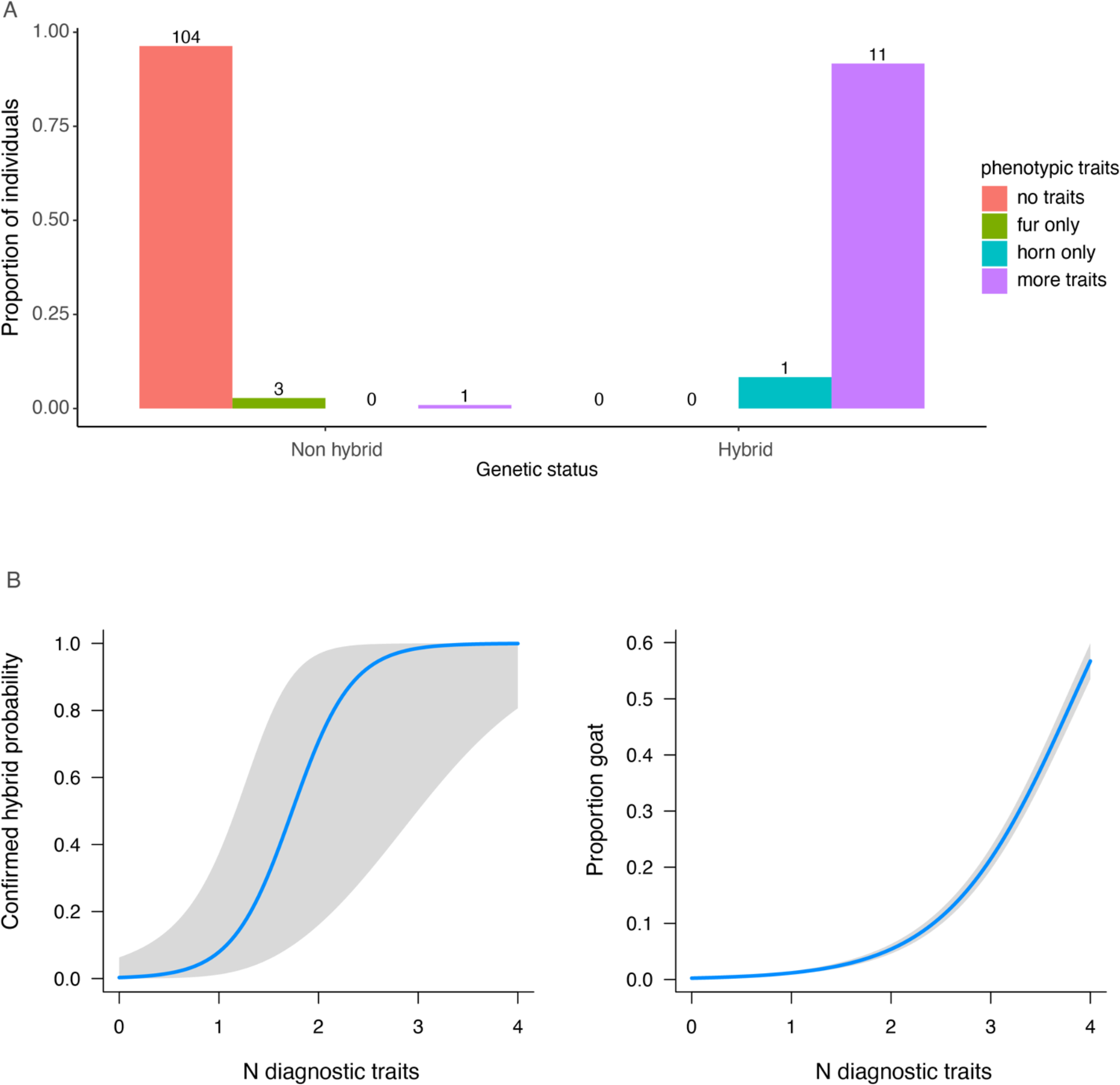
Graphical visualization of the relationship between hybridization and phenotypic diagnostic traits. A) Proportion of individuals showing different diagnostic traits in hybrid and non-hybrid individuals. B) Effect of the number of diagnostic traits on the probability of an individual of being a hybrid. C) Relationship between the observed proportion of goat alleles and the number of diagnostic traits presented by the different individuals.

The number of diagnostic traits was a reliable explanatory variable for the probability of showing some kind of hybridization: logistic regression β=3.34 ± 1.06, *p*=0.002 (Figure 5B). The number of diagnostic traits was also explanatory for the level of hybridization (i.e., the proportion of domestic goat alleles): β=1.57 ± 0.05, *p*<0.001 (Figure 5C).

## Discussion

Our study demonstrates by means of genetic analysis the presence of several hybrids between Alpine ibex and domestic goat. This confirms what was suggested by Moroni et al. (2022) based on phenotypic appearance and also confirms that phenotypic traits can be used to identify recent hybrids between these two species. We also showed the presence of animals with different levels of hybridization (proportion of domestic goat alleles ranging between 0.1 and 0.8), including repeated back-crosses of hybrids in both directions (into Alpine ibex and into domestic goats), suggesting a complete lack of genetic or behavioural reproductive barriers between the two species. Given the low genetic diversity of Alpine ibex, and consequent risk of successful introgression, it is therefore necessary to take management precautions in the areas where domestic goats are sympatric with Alpine ibex to prevent hybridization. This may include higher surveillance of the domestic herds to ensure that all the domestic animals return to the stables at the end of the summer pasture season.

Hybrids were not distributed evenly among the sampling areas. Particularly, we did not find any recent hybrid among the individuals randomly sampled in different populations across the Alps (Italy, France, Switzerland and Austria). The relatively low number of samples collected in those populations may not be enough to completely exclude the presence of hybrids, if they are at low density. However, also Moroni et al. (2022), reported that most of the suspected hybrids were observed in few areas and a previous genetic survey (based on 30 microsatellites) of 1781 Alpine ibex across the Alps also did not detect any recent hybrids except for three known F1 domestic individuals (Grossen et al., 2014). It was hence suggested that hybridisation is rare or hybrids are counter-selected in the wild. Our results instead suggest that hybridisation can be rather strong in particular circumstances as, in the two focal areas, where we genetically confirmed the presence of hybrid swarms (Mayr, 1963). To our knowledge this is the first report of a hybrid swarm between a large mammal and a domesticated relative. In both areas more than one domestic goat was reportedly abandoned in the wild: 3-4 in Gran Paradiso at least 10 years ago (Gran Paradiso National Park Surveillance Service, personal communication) and around 10-15 in Lanzo around 15-20 years ago (Emilio Gugliermetti, personal communication). The feral domestic goats were observed for several years thus increasing mating probabilities with Alpine ibex and further back-crosses. The phenomenon of hybridization in Alpine ibex seems therefore not widely spread at a geographic scale but locally pronounced, also leading, in specific situations (i.e., the release and survival of more than one domestic goat in the same area), to the formation of hybrid swarms.

In the populations where most of the hybrids were identified (Lanzo and Gran Paradiso), the hybrids seemed to be confined to relatively small areas. This is somewhat surprising given that 10 to 20 years have passed since the first abandonment of the domestic goats that likely originated the hybrid swarms. The gregarious behaviour of most *Capra* species as well as philopatry and low dispersal rates typical of Alpine ibex (Brambilla et al., 2020; Marchand et al., 2017; Scillitani et al., 2012) may explain this finding. Although nothing is known about social behaviour of hybrids, in the study areas they have often been observed in mixed groups with other hybrids as well as with Alpine ibex. If gregariousness and philopatry are maintained also in the hybrids, the probability of spread of the domestic goat alleles into other, neighbouring Alpine ibex populations would be less pronounced. However, despite the low dispersal rate, it is known that Alpine ibex, particularly males, may move between sectors (Chauveau et al., 2023.; Marchand et al., 2017), thus constituting possible spreaders for goat alleles, particularly in the Western Alps where the ibex distribution is more continuous (Brambilla et al., 2020).

We observed hybrids of both sex, different age and different generation (Supplementary Table S2) showing that they are able to survive and reproduce and suggesting an absence of behavioural or genetic reproductive barriers. Studies conducted on other mammals reported contrasting results in this respect with some populations struggling to crossbreed, albeit of the same species (red fox, *Vulpes vulpes*, Sacks et al., 2011) and other cases of hybridization between different species (mouflons, *Ovis* spp., Chen et al., 2021) confirming that the potential risk of hybridization and consequent threat for conservation is species-specific. The relatively high number of hybrids observed in the focal areas (we sampled 9 in Lanzo and 3 in Gran Paradiso, but more probable hybrids were observed, at least 20 in Lanzo and 15 in Gran Paradiso, Moroni et al. 2022, and Brambilla personal observation) indicates that at present there are no clear signs of selection against hybrids. In addition, many of the observed male hybrids showed large secondary sexual traits (Figures 3 and 5), suggesting possible hybrid vigour. Alpine ibex are characterized by a polygynous mating system with active defence of oestrous females by dominant males (Brambilla et al., 2022). Hence, large hybrid males may have high reproductive success. Further analysis will be needed for understanding whether both male and female goats and hybrids have the same probability of reproducing.

Although not recent hybrids, all individuals may carry signals of older introgression, which would not be detected by our diagnostic markers as they were specifically designed to identify recent hybridisation (Kessler et al., 2022). Increasing the number of markers let for instance to the reclassification of up to 25% supposedly pure red deer to hybrids, but these were in general more advanced back-crosses (McFarlane et al., 2020). In Alpine ibex, previous genome-wide analyses of introgression based on whole-genome sequencing suggested indeed an average proportion of 2.3% of domestic goat ancestry among non-hybrid Alpine ibex (Münger et al., 2023). This is in the same range as Neanderthal ancestry in humans outside of Africa (1.5-2.1%, Prüfer et al., 2014) and suggests that, although under certain conditions high levels of hybridisation and back-crossing are possible, this phenomenon was generally limited in the past. However, up to present, we cannot conclude whether the general low level of introgression observed in Alpine ibex occurred because of counter-selection of domestic goat alleles in the wild after few generations (which would speak for a general low risk of genetic swamping for this species, similar to what was found in grey wolfs *Canis lupus* in Europe, Pilot et al., 2018) or because less occasions of hybridization occurred in the past and domestic goat alleles were diluted over time by back-crossing into Alpine ibex. Indeed, despite release or abandonment of domestic goat in the wild has likely always happened, several circumstances may have changed in the last decades. First of all, a reduction of good practice in the management of domestic herds may have occurred since, in most cases, domestic goats’ productions (meat or milk) no longer constitute the owners’ primary source of income, which now mostly consists of economic contributions for transhumance to alpine pastures using extensive traditional practices. Second, the increase of Alpine ibex abundance (Brambilla et al., 2020) has enlarged the range of sympatry between the two species and hence the occasions of mating. Expansion of the European wildcat during the last 50 years has also been suggested to be a driving force of introgression from domestic cat, *Felix catus* (Nussberger et al., 2014). Third, the reduction of snow precipitations in the Alps may have reduced winter mortality of both hybrids and domestic goats in the wild, increasing the chances of successful reproduction with Alpine ibex. Winter harshness and snow cover are among the main limiting factor even for wild mountain ungulates adapted to the Alpine climate (Jacobson et al., 2004; Jonas et al., 2008; Rughetti et al., 2011), hence they are likely to have a similar or even stronger effect on feral domestic animals. Milder and dryer winter could instead allow the survival of adult goat and hybrids as well as of new-born, regardless from the birth date (which may occur earlier in the season in hybrids, Giacometti et al., 2004). Climate change has been suggested to be an important player also for hybridisation between mountain hare and the northwards expanding brown hare (Levänen et al., 2018). If the social and environmental changes that occurred in the last decades increase the chances of introgression to occur, hybridization may currently represent a potential threat for the conservation of the species, as it happens for example in Scottish wildcat (Senn et al., 2019) and should be prevented whenever possible. Continued monitoring of the phenomenon of hybridization between Alpine ibex and domestic goat is therefore crucial for a comprehensive understanding of this issue and to enhance the genetic conservation of Alpine ibex.

### Recommendations for hybrid identification and management implications

Our results showed that phenotypic assessment can be a first diagnostic tool to detect hybridization. When more than one phenotypic diagnostic trait was present, the probability of correctly classifying an animal as a hybrid was >80%, going up to 100% with 3 or more traits (Figure 5B). Among the investigated phenotypic traits, the horns seem to be the most reliable, as well as the easiest to identify. All confirmed hybrids had obvious anomalies in the horns (Figure 5A) with a huge variation ranging from complete absence of horns to horns much larger than average. Other anomalies included: triangular section, smoother frontal surface, sometimes also with fusion of the frontal ridges (as typical of some Alpine breed of domestic goat). Pelage colour, instead, especially if not coupled with other traits, does not seem to be a reliable diagnostic trait. Indeed, three animals that showed anomalies in pelage colour (i.e., pelage lighter than common Alpine ibex) but no other diagnostic traits, resulted to be non-hybrids. Those individuals were initially considered as suspected hybrids by local gamekeepers. For one old female of the Lanzo population with two diagnostic traits, instead, the horn length as well as the slightly lighter colour could also be due to the very old age. It is worth to be noted, that none of those four individuals were classified as probable hybrid by the authors of Moroni et al. (2022). Expert-based evaluation of phenotypic appearance could therefore be considered as a relatively reliable diagnostic tool for identification of recent hybrids. For doubtful cases, however, genetic confirmation is the only decisive tool. The genetic classification of hybrids performed in this study relied on 63 diagnostic markers previously designed for this purpose (Kessler et al. 2022). With the requirement of at least two domestic goat alleles, this set of markers would (if missingness is low) allow detecting hybrids up to the 5^th^ generation (or even more if there was mating among hybrids and not just back-crossing into Alpine ibex). Hybrid classification gets however less precise with lower sample quality and higher missingness. In this manuscript we kept samples with a missingness up to 80% and observed, as expected, higher missingness in faecal than tissue samples (Supplementary Figure S1). Collection of faecal samples is a cheap and non-invasive method for DNA collection. However, DNA extracted from faecal samples is often of low quantity and quality (highly fragmented, possible contamination) thus providing less robust results in quantifying the exact proportion of goat alleles (Figure 5). Although we did not conduct a proper analysis to test this, our results suggest that, provided that some precautions are taken, faecal samples can still be used to confirm the status of recent hybrids using the diagnostic markers developed by Kessler et al. (2022), particularly if phenotypic diagnostic traits are present. Faecal samples are instead not recommended if the aim is to know the exact proportion of goat alleles (hence the actual hybrid generation). Precautions that need to be taken when using faecal samples include to freeze the samples as fresh as possible and to repeat at least twice the analysis of each faecal sample and merge reads per individual at the genotyping step. In addition, to further reduce the risk of false positive, we suggest to use stringent genotype filters (minimal genotype quality = 50; minimal read depth = 20), remove individuals with high missingness rate (>80%) and require a hybrid genotype at least at two diagnostic markers to define an individual as a recent hybrid when using faecal samples. With a missingness cut-off of 80% (average of 12 diagnostic markers retained) and the requirement of at least two loci showing a hybrid genotype, we expect that hybrids up to the 3^rd^ generation can be detected, although it would be difficult to determine the specific hybrid category.

Poor management of domestic goat herds grazing in Alpine pastureland led to the creation of hybrid swarms in two Alpine ibex populations. Hybrids are able to survive and reproduce, thus increasing the risk of local genetic swamping and introgression. Given the critical situation of the Alpine ibex gene pool (Biebach & Keller, 2010; Grossen et al., 2018), hybridization may constitute a further threat. Good management practices for domestic animals are therefore necessary to ensure Alpine ibex conservation, particularly in areas where domestic goats are sympatric with Alpine ibex. The same applies to other domestic species coexisting and potentially hybridizing with wild relatives as is the case for domestic cat (Senn et al. 2019), pig (Dzialuk et al., 2018), dog (Pilot et al. 2018), sheep (Šprem et al., 2023) and cattle (Stroupe et al., 2022). When suspected hybrids are present, a systematic survey as described in Moroni et al. (2022) followed by genetic assessment with a method similar to what proposed in this study may constitute a reliable tool to evaluate the abundance of hybrids and their status, to allow sound and informed management decisions.

## Data availability

The data that support the findings of this study are openly available in Dryad Digital Repository at [TBA upon acceptance], reference number [TBA upon acceptance]. Data can be cited as XXXXX et al. [TBA upon acceptance]

## References

Abi-Rached, L., Jobin, M. J., Kulkarni, S., McWhinnie, A., Dalva, K., Gragert, L., Babrzadeh, F., Gharizadeh, B., Luo, M., Plummer, F. A., & others. (2011). The shaping of modern human immune systems by multiregional admixture with archaic humans. Science, 334, 89–94.

Adavoudi, R., & Pilot, M. (2022). Consequences of hybridization in mammals: A systematic review. In Genes (Vol. 13, Issue 1). MDPI. 10.3390/genes13010050

Allendorf, F. W., Leary, R. F., Spruell, P., & Wenburg, J. K. (2001). The problems with hybrids: setting conservation guidelines. TRENDS in Ecology & Evolution, 16(11). http://tree.trends.com0169

Anderson, E. C. (2003). User’s Guide to the Program NewHybrids Version 1.1 beta.

Anderson, E. C., & Thompson, E. A. (2002). A Model-Based Method for Identifying Species Hybrids Using Multilocus Genetic Data. Genetics, 160, 1217–1229. https://academic.oup.com/genetics/article/160/3/1217/6052497

Bickhart, D. M., Rosen, B. D., Koren, S., Sayre, B. L., Hastie, A. R., Chan, S., Lee, J., Lam, E. T., Liachko, I., Sullivan, S. T., Burton, J. N., Huson, H. J., Nystrom, J. C., Kelley, C. M., Hutchison, J. L., Zhou, Y., Sun, J., Crisà, A., Ponce De León, F. A., … Smith, T. P. L. (2017). Single-molecule sequencing and chromatin conformation capture enable de novo reference assembly of the domestic goat genome. Nature Genetics, 49(4), 643–650. 10.1038/ng.3802

Biebach, I., & Keller, L. F. (2010). Inbreeding in reintroduced populations: The effects of early reintroduction history and contemporary processes. Conservation Genetics, 11(2), 527–538. 10.1007/s10592-009-0019-6

Biebach, I., & Keller, L. F. (2012). Genetic variation depends more on admixture than number of founders in reintroduced Alpine ibex populations. Biological Conservation, 147(1), 197–203. 10.1016/j.biocon.2011.12.034

Bolger, A. M., Lohse, M., & Usadel, B. (2014). Trimmomatic: A flexible trimmer for Illumina sequence data. Bioinformatics, 30(15), 2114–2120. 10.1093/bioinformatics/btu170

Bozzuto, C., Biebach, I., Muff, S., Ives, A. R., & Keller, L. F. (2019). Inbreeding reduces long-term growth of Alpine ibex populations. Nature Ecology & Evolution, 3, 1350–1364.

Brambilla, A., Biebach, I., Bassano, B., Bogliani, G., & von Hardenberg, A. (2014). Direct and indirect causal effects of heterozygosity on fitness-related traits in Alpine ibex. Proceedings of the Royal Society B: Biological Sciences, 282(1798). 10.1098/rspb.2014.1873

Brambilla, A., von Hardenberg, A., Canedoli, C., Brivio, F., Sueur, C., & Stanley, C. R. (2022). Long term analysis of social structure: evidence of age-based consistent associations in male Alpine ibex. Oikos, 2022(8). 10.1111/oik.09511

Brambilla, A., Von Hardenberg, A., Nelli, L., & Bassano, B. (2020). Distribution, status, and recent population dynamics of Alpine ibex Capra ibex in Europe. In Mammal Review (Vol. 50, Issue 3, pp. 267–277). Blackwell Publishing Ltd. 10.1111/mam.12194

Canu, A., Vilaça, S. T., Iacolina, L., Apollonio, M., Bertorelle, G., & Scandura, M. (2016). Lack of polymorphism at the MC1R wild-type allele and evidence of domestic allele introgression across European wild boar populations. Mammalian Biology, 81(5), 477–479. 10.1016/j.mambio.2016.01.003

Chauveau, V., Garel, M., Toïgo, C., Anderwald, P., Beurier, M., Bouche, M., Cagnacci, F., Canut, M., Cavailhes, J., Filli, F., Frey-Roos, A., Gressmann, G., Herfindal, I., Martinelli, L., Papet, R., Petit, E., Ramanzin, M., Vannard, E., Loison, A., … Marchand, P. (n.d.). Identifying the environmental drivers of corridors and predicting 1 connectivity between seasonal ranges in multiple populations of Alpine ibex 2 (Capra ibex) as tools for conserving migration. 10.1101/2023.03.02.530594

Chen, Z. H., Xu, Y. X., Xie, X. L., Wang, D. F., Aguilar-Gómez, D., Liu, G. J., Li, X., Esmailizadeh, A., Rezaei, V., Kantanen, J., Ammosov, I., Nosrati, M., Periasamy, K., Coltman, D. W., Lenstra, J. A., Nielsen, R., & Li, M. H. (2021). Whole-genome sequence analysis unveils different origins of European and Asiatic mouflon and domestication-related genes in sheep. Communications Biology, 4(1). 10.1038/s42003-021-02817-4

Couturier, M. A. (1962). Le Bouquetin des Alpes: capra aegagrus ibex ibex L. Couturier.

Danecek, P., Auton, A., Abecasis, G., Albers, C. A., Banks, E., DePristo, M. A., Handsaker, R. E., Lunter, G., Marth, G. T., Sherry, S. T., McVean, G., & Durbin, R. (2011). The variant call format and VCFtools. Bioinformatics, 27(15), 2156–2158. 10.1093/bioinformatics/btr330

Dzialuk, A., Zastempowska, E., Skórzewski, R., Twarużek, M., & Grajewski, J. (2018). High domestic pig contribution to the local gene pool of free-living European wild boar: a case study in Poland. Mammal Research, 63(1), 65–71. 10.1007/s13364-017-0331-3

Ferguson, M. M., Danzmann, R. G., & Allendorf, F. W. (1988). Developmental success of hybrids between two taxa of salmonid fishes with moderate structural gene divergence. Canadian Journal of Zoology, 66(6), 1389–1395. 10.1139/z88-204

Fukui, S., May-McNally, S. L., Taylor, E. B., & Koizumi, I. (2018). Maladaptive secondary sexual characteristics reduce the reproductive success of hybrids between native and non-native salmonids. Ecology and Evolution, 8(23), 12173–12182. 10.1002/ece3.4676

Genovart, M. (2009). Natural hybridization and conservation. Biodiversity and Conservation, 18(6), 1435–1439. 10.1007/s10531-008-9550-x

Giacometti, M., Roganti, R., De Tann, D., Stahlberger-Saitbekova, N., & Obexer-Ruff, G. (2004). Alpine ibex Capra ibex ibex x domestic goat C. aegagrus domestica hybrids in a restricted area of southern Switzerland. Wildlife Biology, 10(2), 137–143. 10.2981/wlb.2004.018

Gouy, A., Excoffier, L., & Nielsen, R. (2020). Polygenic patterns of adaptive introgression in modern humans are mainly shaped by response to pathogens. Molecular Biology and Evolution, 37(5), 1420–1433. 10.1093/molbev/msz306

Grossen, C., Biebach, I., Angelone-Alasaad, S., Keller, L. F., & Croll, D. (2018). Population genomics analyses of European ibex species show lower diversity and higher inbreeding in reintroduced populations. Evolutionary Applications, 11(2), 123–139. 10.1111/eva.12490

Grossen, C., Guillaume, F., Keller, L. F., & Croll, D. (2020). Purging of highly deleterious mutations through severe bottlenecks in Alpine ibex. Nature Communications, 11(1). 10.1038/s41467-020-14803-1

Grossen, C., Keller, L., Biebach, I., Zhang, W., Tosser-Klopp, G., Ajmone, P., Amills, M., Boitard, S., Chen, W., Cheng, S., Dong, Y., Faraut, T., Faruque, O., Heuven, H., Jinshan, Z., Jun, L., Lenstra, H., Li, X., Liu, X., … Croll, D. (2014). Introgression from Domestic Goat Generated Variation at the Major Histocompatibility Complex of Alpine Ibex. PLoS Genetics, 10(6). 10.1371/journal.pgen.1004438

Harbicht, A., Wilson, C. C., & Fraser, D. J. (2014). Does human-induced hybridization have long-term genetic effects? Empirical testing with domesticated, wild and hybridized fish populations. Evolutionary Applications, 7(10), 1180–1191. 10.1111/eva.12199

Howard-McCombe, J., Jamieson, A., Carmagnini, A., Russo, I.-R. M., Ghazali, M., Campbell, R., Driscoll, C., Murphy, W. J., Nowak, C., O’Connor, T., Tomsett, L., Lyons, L. A., Muñoz-Fuentes, V., Bruford, M. W., Kitchener, A. C., Larson, G., Frantz, L., Senn, H., Lawson, D. J., & Beaumont, M. A. (2023). Genetic swamping of the critically endangered Scottish wildcat was recent and accelerated by disease. Current Biology: CB, 33(21), 4761–4769.e5. 10.1016/j.cub.2023.10.026

Iacolina, L., Corlatti, L., Buzan, E., Safner, T., & Šprem, N. (2019). Hybridisation in European ungulates: an overview of the current status, causes, and consequences. In Mammal Review (Vol. 49, Issue 1, pp. 45–59). Blackwell Publishing Ltd. 10.1111/mam.12140

Jacobson, A. R., Provenzale, A., Von Hardenberg, A., Bassano, B., & Festa-Bianchet, M. (2004). Climate forcing and density dependence in a mountain ungulate population. Ecology, 85(6), 1598–1610.

Jonas, T., Geiger, F., & Jenny, H. (2008). Mortality pattern of the Alpine chamois: The influence of snow-meteorological factors. Annals of Glaciology, 49, 56–62. 10.3189/172756408787814735

Kessler, C., Brambilla, A., Waldvogel, D., Camenisch, G., Biebach, I., Leigh, D. M., Grossen, C., & Croll, D. (2022). A robust sequencing assay of a thousand amplicons for the high-throughput population monitoring of Alpine ibex immunogenetics. Molecular Ecology Resources, 22(1), 66–85. 10.1111/1755-0998.13452

Levänen, R., Thulin, C. G., Spong, G., & Pohjoismäki, J. L. O. (2018). Widespread introgression of mountain hare genes into Fennoscandian brown hare populations. PLoS ONE, 13(1). 10.1371/journal.pone.0191790

Li, H. (2013). Aligning sequence reads, clone sequences and assembly contigs with BWA-MEM. http://arxiv.org/abs/1303.3997

Lischer, H. E. L., & Excoffier, L. (2012). PGDSpider: An automated data conversion tool for connecting population genetics and genomics programs. Bioinformatics, 28(2), 298–299. 10.1093/bioinformatics/btr642

Magoč, T., & Salzberg, S. L. (2011). FLASH: Fast length adjustment of short reads to improve genome assemblies. Bioinformatics, 27(21), 2957–2963. 10.1093/bioinformatics/btr507

Marchand, P., Freycon, P., Herbaux, J. P., Game, Y., Toïgo, C., Gilot-Fromont, E., Rossi, S., & Hars, J. (2017). Sociospatial structure explains marked variation in brucellosis seroprevalence in an Alpine ibex population. Scientific Reports, 7(1). 10.1038/s41598-017-15803-w

Mayr, E. (1963). Animal species and evolution. Harvard University Press.

McFarlane, S. E., Hunter, D. C., Senn, H. V., Smith, S. L., Holland, R., Huisman, J., & Pemberton, J. M. (2020). Increased genetic marker density reveals high levels of admixture between red deer and introduced Japanese sika in Kintyre, Scotland. Evolutionary Applications, 13(2), 432–441. 10.1111/eva.12880

McFarlane, S. E., & Pemberton, J. M. (2019). Detecting the True Extent of Introgression during Anthropogenic Hybridization. In Trends in Ecology and Evolution (Vol. 34, Issue 4, pp. 315– 326). Elsevier Ltd. 10.1016/j.tree.2018.12.013

Moroni, B., Brambilla, A., Rossi, L., Meneguz, P. G., Bassano, B., & Tizzani, P. (2022). Hybridization between Alpine Ibex and Domestic Goat in the Alps: A Sporadic and Localized Phenomenon? Animals, 12(6). 10.3390/ani12060751

Münger, X. C. T., & Grossen, C. (2020). Genome-wide analysis of introgression from domestic to a wild goat.

Münger, X., Robin, M., Dalén, L., & Grossen, C. (2023). Facilitated introgression from domestic goat into Alpine ibex at immune loci. BioRxiv: 568345. 10.1101/2023.11.23.568345

Negri, A., Pellegrino, I., Mucci, N., Randi, E., Tizzani, P., Meneguz, P. G., & Malacarne, G. (2013). Mitochondrial DNA and microsatellite markers evidence a different pattern of hybridization in red-legged partridge (Alectoris rufa) populations from NW Italy. European Journal of Wildlife Research, 59(3), 407–419. 10.1007/s10344-012-0686-3

Nussberger, B., Wandeler, P., Weber, D., & Keller, L. F. (2014). Monitoring introgression in European wildcats in the Swiss Jura. Conservation Genetics, 15(5), 1219–1230. 10.1007/s10592-014-0613-0

Pidancier, N., Jordan, S., Luikart, G., & Taberlet, P. (2006). Evolutionary history of the genus Capra (Mammalia, Artiodactyla): Discordance between mitochondrial DNA and Y-chromosome phylogenies. Molecular Phylogenetics and Evolution, 40(3), 739–749. 10.1016/j.ympev.2006.04.002

Pilot, M., Greco, C., vonHoldt, B. M., Randi, E., Jędrzejewski, W., Sidorovich, V. E., Konopiński, M. K., Ostrander, E. A., & Wayne, R. K. (2018). Widespread, long-term admixture between grey wolves and domestic dogs across Eurasia and its implications for the conservation status of hybrids. Evolutionary Applications, 11(5), 662–680. 10.1111/eva.12595

Pohjoismäki, J. L. O., Michell, C., Levänen, R., & Smith, S. (2021). Hybridization with mountain hares increases the functional allelic repertoire in brown hares. Scientific Reports, 11(1). 10.1038/s41598-021-95357-0

Prüfer, K., Racimo, F., Patterson, N., Jay, F., Sankararaman, S., Sawyer, S., Heinze, A., Renaud, G., Sudmant, P. H., De Filippo, C., Li, H., Mallick, S., Dannemann, M., Fu, Q., Kircher, M., Kuhlwilm, M., Lachmann, M., Meyer, M., Ongyerth, M., … Pääbo, S. (2014). The complete genome sequence of a Neanderthal from the Altai Mountains. Nature, 505(7481), 43–49. 10.1038/nature12886

R Core Team. (2022). R: A language and environment for statistical computing. R Foundation for Statistical Computing. https://www.R-project.org/

Randi, E. (2008). Detecting hybridization between wild species and their domesticated relatives. In Molecular Ecology (Vol. 17, Issue 1, pp. 285–293). 10.1111/j.1365-294X.2007.03417.x

Rhymer, J. M., & Simberloff, D. (1996). Extinction by hybridization and introgression. In Annu. Rev. Ecol. Syst (Vol. 27). www.annualreviews.org

Rughetti, M., Toïgo, C., Von Hardenberg, A., Rocchia, E., & Festa-Bianchet, M. (2011). Effects of an exceptionally snowy winter on chamois survival. Acta Theriologica, 56(4), 329–333. 10.1007/s13364-011-0040-2

Sacks, B. N., Moore, M., Statham, M. J., & Wittmer, H. U. (2011). A restricted hybrid zone between native and introduced red fox (Vulpes vulpes) populations suggests reproductive barriers and competitive exclusion. Molecular Ecology, 20(2), 326–341. 10.1111/j.1365-294X.2010.04943.x

Scillitani, L., Sturaro, E., Menzano, A., Rossi, L., Viale, C., & Ramanzin, M. (2012). Post-release spatial and social behaviour of translocated male Alpine ibexes (Capra ibex ibex) in the eastern Italian Alps. European Journal of Wildlife Research, 58(2), 461–472. 10.1007/s10344-011-0596-9

Senn, H. V., Ghazali, M., Kaden, J., Barclay, D., Harrower, B., Campbell, R. D., Macdonald, D. W., & Kitchener, A. C. (2019). Distinguishing the victim from the threat: SNP-based methods reveal the extent of introgressive hybridization between wildcats and domestic cats in Scotland and inform future in situ and ex situ management options for species restoration. Evolutionary Applications, 12(3), 399–414. 10.1111/eva.12720

Simberloff, D. (1996). Hybridization between native and introduced wildlife species: Importance for conservation. Wildlife Biology, 2(3), 143–150. 10.2981/wlb.1996.012

Skaala, Ø., Besnier, F., Borgstrøm, R., Barlaup, B. T., Sørvik, A. G., Normann, E., Østebø, B. I., Hansen, M. M., & Glover, K. A. (2019). An extensive common-garden study with domesticated and wild Atlantic salmon in the wild reveals impact on smolt production and shifts in fitness traits. Evolutionary Applications, 12(5), 1001–1016. 10.1111/eva.12777

Šprem, N., Buzan, E., & Safner, T. (2023). How we look: European wild mouflon and feral domestic sheep hybrids. Current Zoology. 10.1093/cz/zoad031

Stroupe, S., Forgacs, D., Harris, A., Derr, J. N., & Davis, B. W. (2022). Genomic evaluation of hybridization in historic and modern North American Bison (Bison bison). Scientific Reports, 12(1). 10.1038/s41598-022-09828-z

Stüwe, M., & Nievergelt, B. (1991). Recovery of Alpine ibex from near extinction: the result of effective protection, captive breeding, and reintroductions. Applied Animal Behaviour Science, 29(1–4), 379–387.

Toïgo, C., Brambilla, A., Grignolio, S., & Pedrotti, L. (2020). Capra ibex. The IUCN Red List of Threatened Species. The IUCN Red List of Threatened Species.

