## Supplementary for "Genetic evidence of a hybrid swarm between Alpine ibex and domestic goat"

### Supplementary Figures

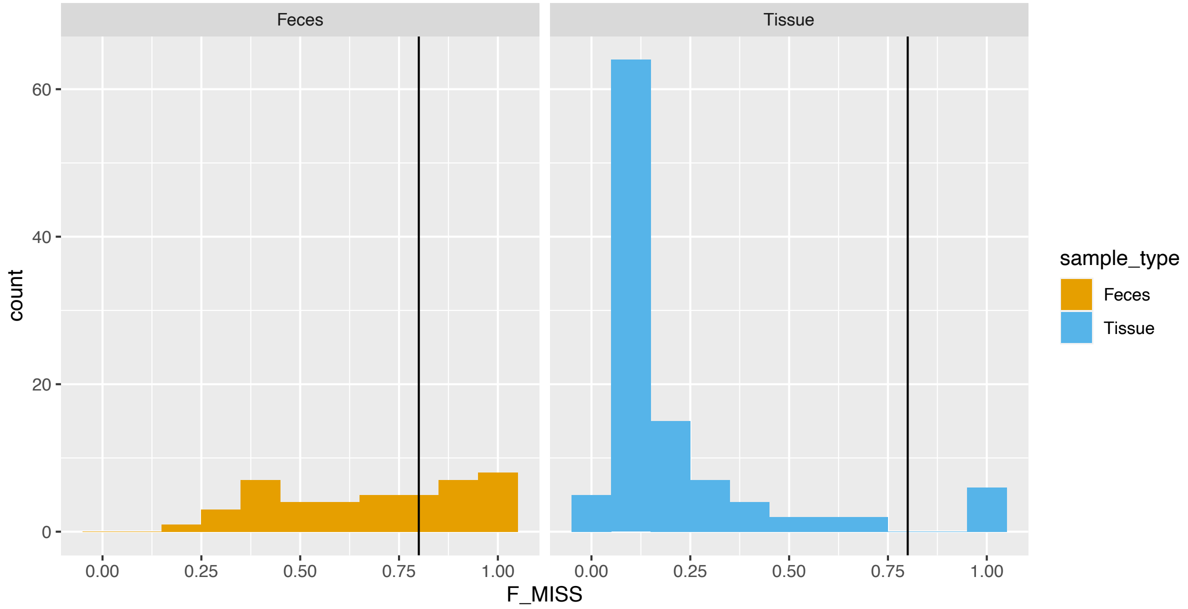


**Supplementary Figure S1:** Number of samples of different kind (Faeces in yellow and Tissue in light blue) with different proportion of missingness. Black line represents the threshold used in this study to filter out samples with high missingness (80%).

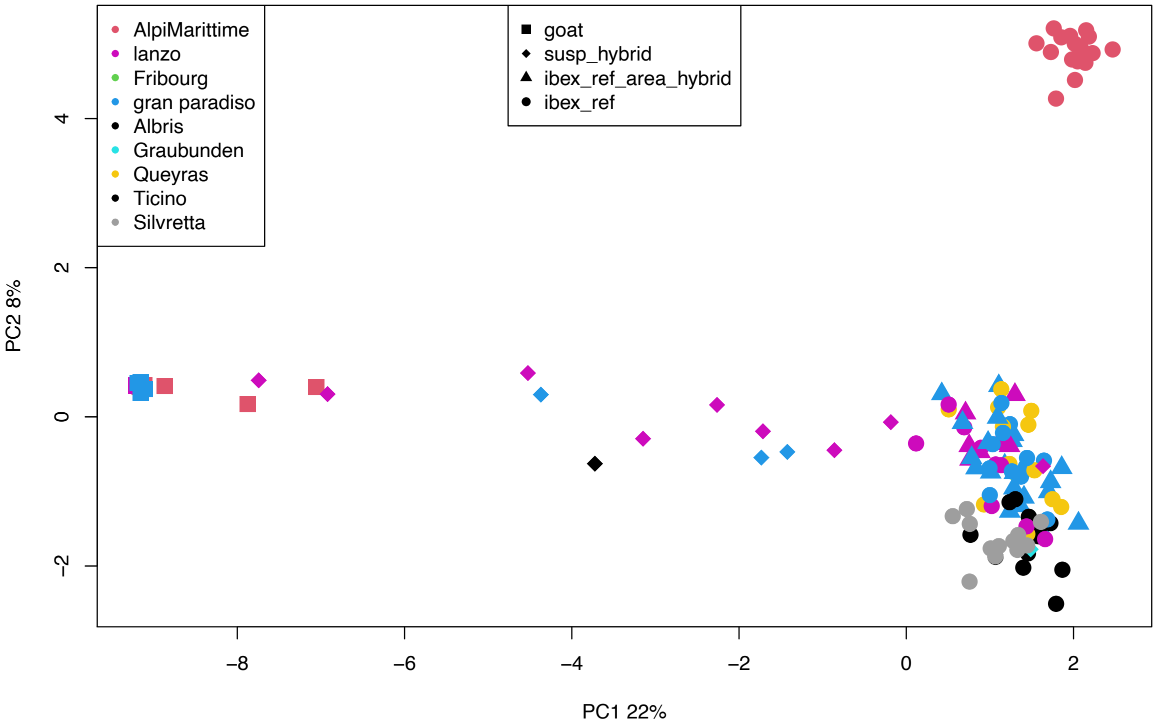

**Supplementary Figure S2:** Principal component analysis (PCA) based on 465 neutral SNPs on 133 individuals. Colors indicate different Alpine ibex populations. Shape indicate the category of the individuals included (dot: non-suspected Alpine ibex outside focal area (ibex_ref_area_hybrid), triangle: non-suspected Alpine ibex focal area (ibex_ref), rhombus: suspected hybrids (susp_hybrid) and square: non-suspected domestic goats (goat).

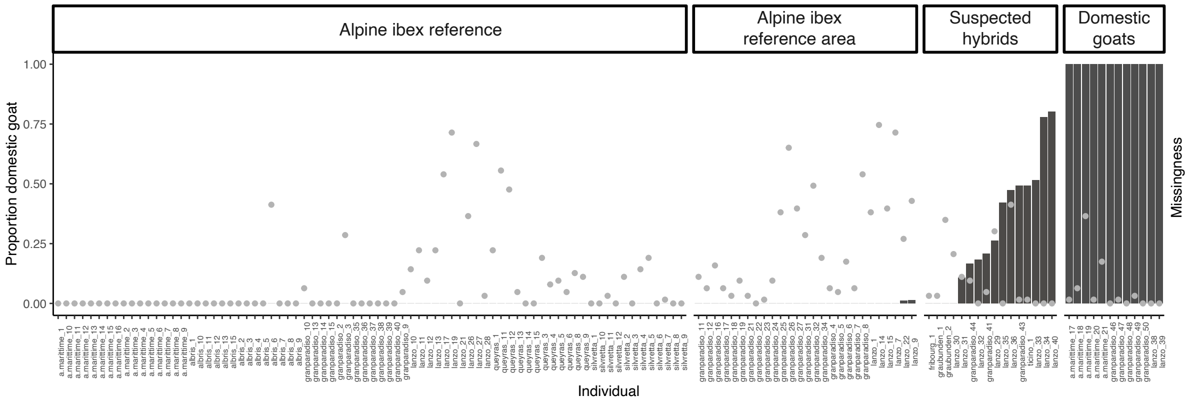


**Supplementary Figure S3:** Proportion domestic goat ancestry (based on 63 diagnostic SNPs, shown in bars) and individual missingness (grey dots).

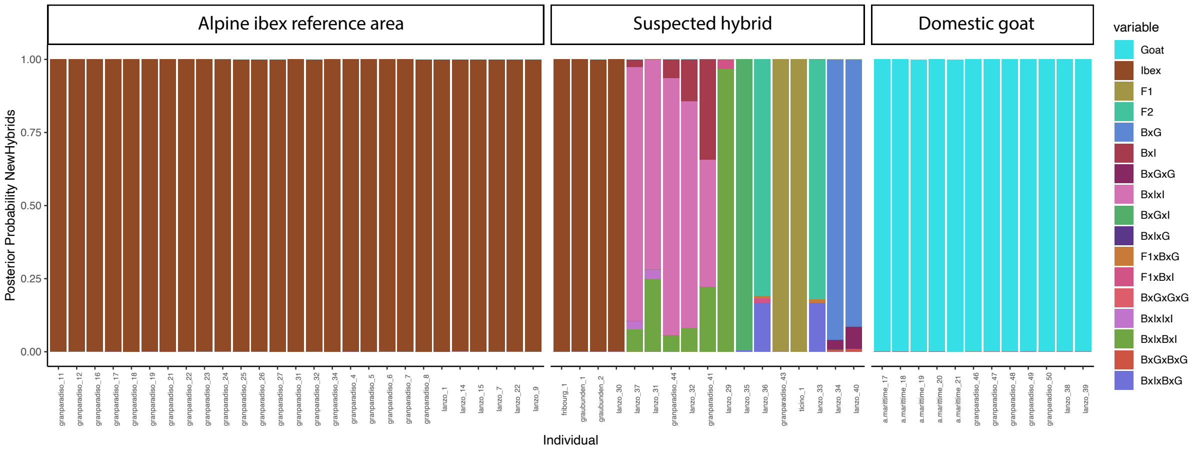


**Supplementary Figure S4:** Cumulative probabilities computed by NewHybrids for each individual to belong to one of the pre- defined hybrid classes. The plot includes visualization of all tested categories. Categories likely occurring at the same generation were merged in the main figure (Fig, 2A) to improve readability. See Table S1 for the custom genotype frequencies used for each category in the NewHybrids analysis.

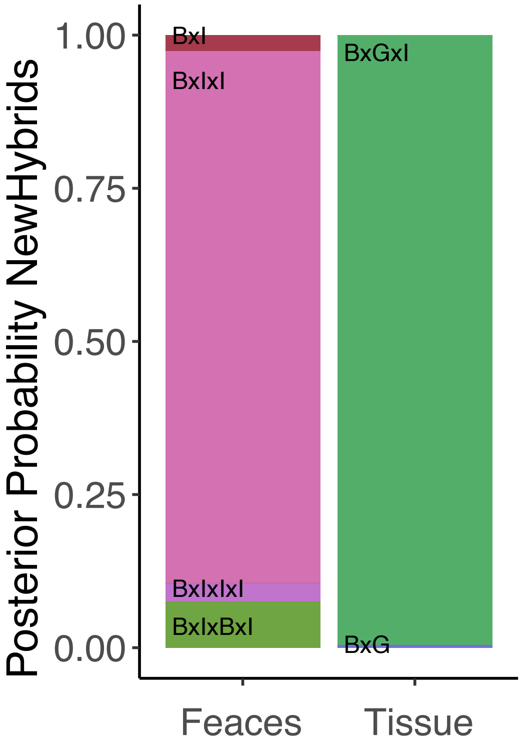


**Supplementary Figure S5:** Cumulative probabilities computed by NewHybrids for the two samples (tissue and faecal) collected from the same individual to belong to one of the pre-defined hybrid classes. The plot includes visualization of all tested categories.

#### Supplementary Tables

**Table S1:** Custom genotype frequencies table used for NewHybrids analysis.

| | | | | |
|---|---|---|---|---|
| 17 | | | | |
| Pure_G | 1.00000000 | 0.00000000 | 0.00000000 | 0.00000000 |
| Pure_I | 0.00000000 | 0.00000000 | 0.00000000 | 1.00000000 |
| F1 | 0.00000000 | 0.50000000 | 0.50000000 | 0.00000000 |
| F2 | 0.25000000 | 0.25000000 | 0.25000000 | 0.25000000 |
| BxG | 0.50000000 | 0.25000000 | 0.25000000 | 0.00000000 |
| BxI | 0.00000000 | 0.25000000 | 0.25000000 | 0.50000000 |
| BxGxG | 0.75000000 | 0.12500000 | 0.12500000 | 0.00000000 |
| BxIxI | 0.00000000 | 0.12500000 | 0.12500000 | 0.75000000 |
| BxGxI | 0.00000000 | 0.37500000 | 0.37500000 | 0.25000000 |
| BxIxG | 0.25000000 | 0.37500000 | 0.37500000 | 0.00000000 |
| F1xBxG | 0.37500000 | 0.25000000 | 0.25000000 | 0.12500000 |
| F1xBxI | 0.12500000 | 0.25000000 | 0.25000000 | 0.37500000 |
| BxGxGxG | 0.87500000 | 0.06250000 | 0.06250000 | 0.00000000 |
| BxIxIxI | 0.00000000 | 0.06250000 | 0.06250000 | 0.87500000 |
| BxIxBxI | 0.06250000 | 0.18750000 | 0.18750000 | 0.56250000 |

| | | | | |
|---|---|---|---|---|
| BxGxBxG | 0.56250000 | 0.18750000 | 0.18750000 | 0.06250000 |
| BxIxBxG | 0.18750000 | 0.31250000 | 0.31250000 | 0.18750000 |
| Pure_G | 1.00000000 | 0.00000000 | 0.00000000 | 0.00000000 |

**Table S2:** List of all samples collected from suspected hybrids. The table includes sample id, sample type, population, sex, number of missing loci and the proportion of goat alleles (P goat) for the samples that passed the missingness filter. Last column shows which individuals were born in captivity. Samples are ordered by sample type and missingness.

| Sample id | Sample type | Population | Sex | N missing loci | P goat | Note ind. |
|---|---|---|---|---|---|---|
| valsusa_01 | FTA card | | f | 63 | - | Domestic |
| queyras_16 | Feces | Val Varaita (Queyras) | m | | - | |
| rocciamelone_1 | Feces | Val Susa (Rocciamelone) | m | 63 | - | |
| lanzo_31 | Feces | Lanzo | m | 7 | 0.11 | |
| lanzo_30 | Feces | Lanzo | f | 13 | 0.00 | |
| lanzo_29 | Feces | Lanzo | f | 19 | 0.26 | |
| lanzo_36 | Feces | Lanzo | f | 26 | 0.47 | |
| lanzo_37 | Feces | Lanzo | m | 44 | 0.11 | |
| lanzo_40 | Tissue | | f | 0 | 0.80 | Domestic |
| lanzo_32 | Tissue | Lanzo | m | 0 | 0.18 | |
| lanzo_33 | Tissue | Lanzo | f | 0 | 0.52 | |
| lanzo_34 | Tissue | Lanzo | f | 0 | 0.78 | |
| lanzo_35 | Tissue | Lanzo | m | 0 | 0.42 | |
| ticino_1 | Tissue | | f | 1 | 0.49 | Domestic |
| granparadiso_43 | Tissue | Gran Paradiso | m | 1 | 0.49 | |
| graubunden_1 | Tissue | Graubunden | NA | 2 | 0.00 | |
| fribourg_1 | Tissue | Fribourg | m | 2 | 0.00 | |
| granparadiso_41 | Tissue | Gran Paradiso | m | 3 | 0.21 | |
| granparadiso_44 | Tissue | Gran Paradiso | m | 6 | 0.17 | |
| graubunden_2 | Tissue | Graubunden | NA | 22 | 0.00 | |
